# Exploring the Architectural Biases of the Canonical Cortical Microcircuit

**DOI:** 10.1101/2024.05.23.595629

**Authors:** Aishwarya Balwani, Suhee Cho, Hannah Choi

## Abstract

The cortex plays a crucial role in various perceptual and cognitive functions, driven by its basic unit, the *canonical cortical microcircuit*. Yet, we remain short of a framework that definitively explains the structure-function relationships of this fundamental neuroanatomical motif. To better understand how physical substrates of cortical circuitry facilitate their neuronal dynamics, we employ a computational approach using recurrent neural networks and representational analyses. We examine the differences manifested by the inclusion and exclusion of biologically-motivated inter-areal laminar connections on the computational roles of different neuronal populations in the microcircuit of two hierarchically-related areas, throughout learning. Our findings show that the presence of feedback connections correlates with the functional modularization of cortical populations in different layers, and provides the microcircuit with a natural inductive bias to differentiate expected and unexpected inputs at initialization. Furthermore, when testing the effects of training the microcircuit and its variants with a predictive-coding inspired strategy, we find that doing so helps better encode noisy stimuli in areas of the cortex that receive feedback, all of which combine to suggest evidence for a predictive-coding mechanism serving as an intrinsic operative logic in the cortex.

## 1 Introduction

The brain is known to extensively implement hierarchical, predictive computations for both learning and inference. In partricular, predictive coding theory (1; 2) posits that the cortex relies on the differences arising between top-down predictions (made by areas higher in the hierarchy), and bottom-up sensory information (received by areas lower in the hierarchy) for efficient information processing and perception (3; 4; 5; 6). Numerous experimental studies also support this theory, e.g., when trained to perform a visual change detection task or when learning a sequence of images, mice have been observed to exhibit heightened neural activity when viewing “surprise” inputs, with their neuronal representations varying appreciably in different cortical layers and areas depending on whether the stimuli are expected or unexpected (7; 8). Additionally, unexpected event signals have also been shown to predict subsequent changes in responses to expected and unexpected stimuli in both individual (9) and populations of neurons (10), making a case for the neocortex indeed instantiating a predictive hierarchical model wherein unexpected events and error signals drive learning.

The anatomical substrate for these computations is the **canonical cortical microcircuit** (11; 12; 13), a structural motif that is shared across cortical areas and found in multiple mammalian species (14; 15; 16). Spanning the length of the cortical column, the microcircuit is defined by distinctive intra- and inter-areal layer-specific connectivity patterns; specifically feedforward signals stemming from the shallow layers (2/3) of the lower cortical area and targeting the granular layer (4) of the higher area, alongwith feedback signals that arise from the deep layers (5/6) of the higher area, and project onto layers (2/3) and (5/6) of the lower area. In connection with the predictive coding theory, these distinct layer-specific projection patterns are postulated to shape computations of expectations and surprisal along the cortical hierarchy. Furthermore, it is also suggested that layers receiving feedback (i.e., 2/3 and 5/6) from higher areas counter the driving feedforward signals originating from superficial layers (2/3), and are therefore the location at which the error/difference computations that are crucial for the predictive coding hypothesis to hold, occur (17; 18).

Indeed, previous studies have provided indirect experimental evidence for prediction-related computations in various cortical layers and areas (19; 20; 21), as well as considering in an abstract sense the theoretical underpinnings of predictive coding (22; 23), their normative formulations (24), and how they might arise in a Bayesian framework (25; 26). However, current understanding of how the structural and computational primitives of the cortical microcircuit shape its representations to induce relevant encoding of errors and (un)expectedness of stimuli within it is limited (27; 28). Given the current state of technology, *in vivo* efforts to experimentally test the underlying mechanisms of predictive coding still face enormous challenges; We therefore propose to explore these questions *in silico*, building on a body of work where deep neural networks have been successfully utilized as models of various neurobiological systems(29; 30; 31; 32; 33; 34). Employing an anatomically-informed ensemble of recurrent neural networks (RNNs) to represent the canonical cortical microcircuit, we interrogate the following aspects of its role in learning: 1) Where are task-related variables encoded across the microcircuit? How do inter-areal feedback connections shape these neural representations? 2) Are there inherent performance advantages associated with the canonical architectural motif? And 3) Are the geometries and functionalities of layer-specific neural representations primarily influenced by the architecture itself, or more so by the training scheme used when learning the task?

By studying representations from multiple architectures with systematically altered structural motifs across different sequential learning tasks, we find that the inherent architectural hierarchy of the microcircuit naturally leads to differences in the geometries of representations across the cortical hierarchy. We also observe that the combination of biologically-consistent inter-areal connections and time-delays in signal projections provide an intrinsic inductive bias for the segregation of simple expected and unexpected stimuli at initialization in layers that receive inter-areal feedback. Additionally, there is an emergence of further functional specialization of neuronal populations in terms of representing task-relevant variables over training. Finally, the inclusion of a predictive-coding based training strategy further strengthens this modularization, specifically in the context of encoding unexpectedness in the stimuli. Taken together, these results suggest a fundamental predictive-coding based mechanism at work in the cortex.

## 2 Models and Methods

In this section we describe the canonical cortical microcircuit as observed *in vivo*, followed by the details of our anatomically-constrained model *in silico*. We subsequently explain the structural modifications we make to our model as to study the effects of inter-areal feedback on learning and performance in the microcircuit. We also provide brief descriptions of the tasks we train our models on, and the methods we use to analyze the representations across the circuit across learning.

### 2.1 The Canonical Cortical Microcircuit: *In-Vivo & In-Silico*

#### Standard *in-vivo* motif

Building on previous studies (15; 14; 16), following are the “universal” set of structural connections upon which our anatomically-constrained, *in silico*, system is modelled:i) Feedforward inputs are received at Layer 4, which then projects to superficial Layers 2/3, ii) Layers 2/3 project both within and across the column, the former set of connections being directed to the deeper Layers 5/6 (35), and the latter being sent as feedforward signal to the next higher order area’s Layer 4 (36), and iii) Layers 5/6 subsequently also project both within and across the column, the former being directed to Layer 4, and the latter set of connections acting inter-areal feedback to the previous lower area’s Layers 2/3 and Layers 5/6.

#### Standard *in-silico* motif

Our typical model, which we call the corticalRNN (Fig. 1.A), is comprised of a lower and higher processing area (or column), each of which utilizes an ensemble of three Elman RNNs (37). The internal state (*h*^(*t*)^) of each RNN at time *t* is determined as:

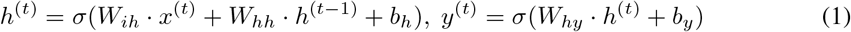

where *x*^(*t*)^ are the inputs received by the RNN at time *t, W*_*ih*_ is the input channel that projects *x*^(*t*)^ onto the RNN’s neuronal population, *W*_*hh*_ are the recurrent weights that specify the connectivity of neurons within the RNN, *h*^(*t*−1)^ is the internal state of the network from the previous time-step, and *b*_*h*_ is a bias term. *y*^(*t*)^ is subsequently the output or readout of the RNN, *W*_*hy*_ projects the current hidden state *h*_*t*_ onto the appropriate output space, and *b*_*y*_ is another bias term. *σ* is a non-linear activation function, e.g., tanh. Every individual RNN models the dynamics of one of the cortical neuronal populations in superficial Layers 2/3, granular Layer 4, or deep Layers 5/6.

**Figure 1:**
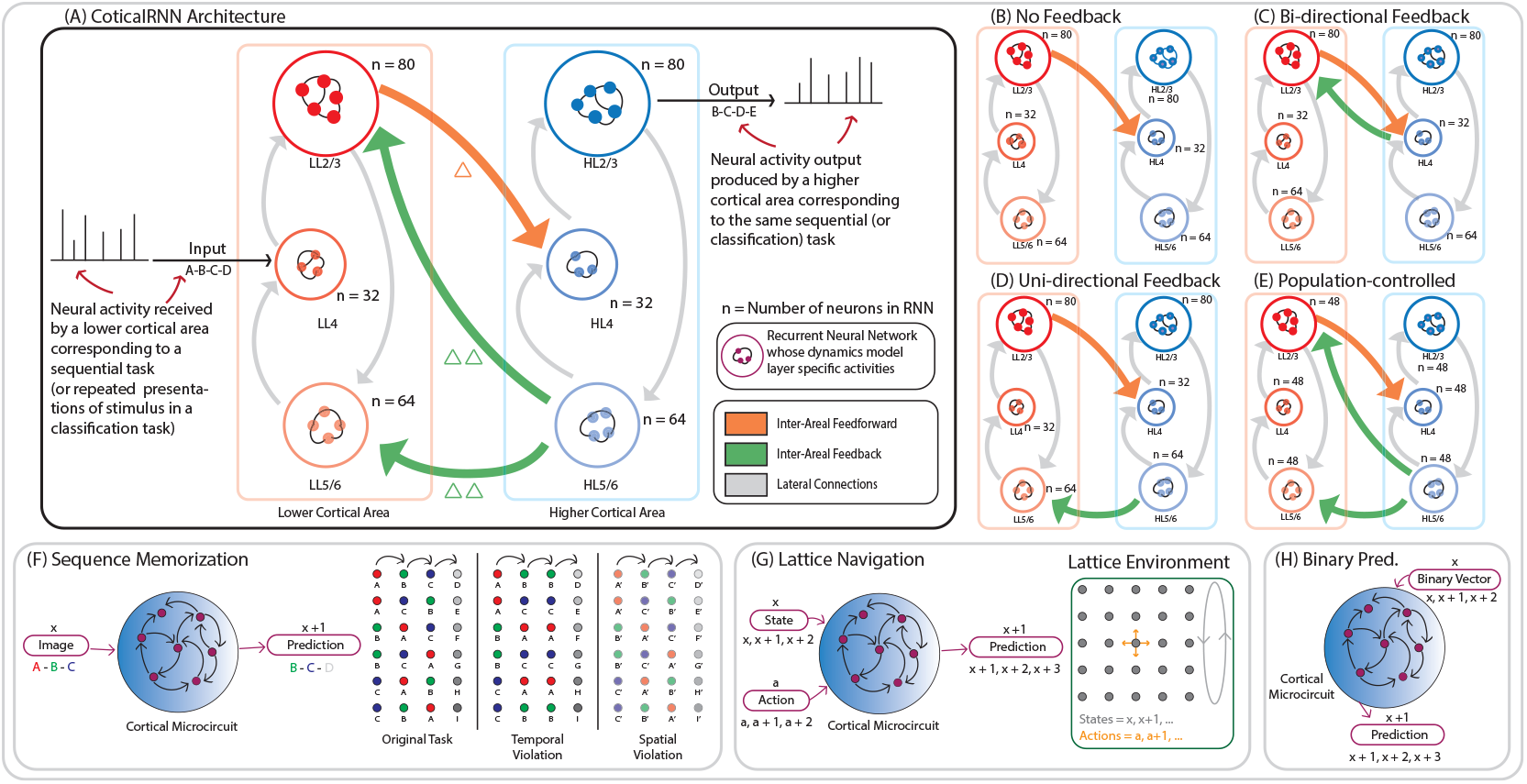
Experimental Setup. (A) Architecture of the corticalRNN. Various arrows show the direction of connectivity across the microcircuit, while each Δ represents one time-step of delay. (B)-(E) Structurally altered versions of the corticalRNN motif. (F)-(H) Schematics of our sequential learning tasks, along with (F) Graphical representations of our temporal and spatial perturbations.

Inter-areal connectivity – i.e., feedforward and feedback projections – as well as the intra-areal connectivity (i.e., lateral projections) between the various RNNs representing cortical layers are modelled by learnable projection matrices with fixed, sparse connectivity patterns sampled from a Bernoulli distribution. The directionalities of both intra and inter-areal connectivity follow the descriptions laid out previously, while the relative sizes of the neuronal populations in the RNNs roughly correspond to the specifications established in the literature with respect to the mouse neuroanatomy, specifically the visual cortex (38). To that end, projections from lateral (i.e., within an area or column) and inter-areal feedback are incorporated by a recipient RNN after a delay of one and two time-steps respectively. All long-range inter-areal connections are strictly excitatory.

The data received and processed at every layer of the canonical microcircuit are as follows:

- **Lower area layers: LL4** : *input* + *proj*(*LL* 5*/*6 → *LL* 4)^4^, **LL2/3** : *proj*(*LL* 4 → *LL* 2*/*3) + *FB*_*a*_, **LL5/6** : *proj*(*LL* 2*/*3 → *LL* 5*/*6) + *FB*_*b*_
- **Higher area layers: HL4** : *FF* + *proj*(*HL* 5*/*6 → *HL* 4), **HL2/3** : *proj*(*HL* 4 → *HL* 2*/*3), **HL5/6** : *proj*(*HL* 2*/*3 → *HL* 5*/*6)

Here, *proj*(*a* → *b*) represents the projection of activity from any layer *a* onto layer *b*. The feedforward projection *FF* is defined as *proj*(*LL*2*/*3 → *HL*4). The two feedback projections *FB*_*a*_, *FB*_*b*_ are defined as *proj*(*HL*5*/*6 → *LL*2*/*3) and *proj*(*HL*5*/*6 → *LL*5*/*6) respectively. Explicit computations carried out by each of the layers are provided in Appendix A.

#### Structurally altered motifs

To isolate the effects of the canonical inter-areal feedback connections on learning and performance in the microcircuit, we study the following variations of the motif: (i) **No inter-areal feedback** from the higher to the lower cortical area, i.e., LL2/3 ↚ HL5/6, and LL5/6 ↚ HL5/6 (Fig. 1.B), (ii) A single, weighted, **bi-directional feedforward-feedback connection** between the lower and higher cortical areas (LL2/3 ⇌ HL4) (Fig. 1.C), (iii) **Uni-directional feedback**, where the connection LL5/6 ← HL5/6 is preserved while the connection LL2/3 ↚ HL5/6 (which in the canonical architecture could interact with the feedforward projection at LL2/3), is removed (Fig. 1.D), (iv) The skeleton of the microcircuit is preserved, but number of neurons across all RNNs is equalized, i.e., **same-sized populations**, doing away with any physical compression and expansion of signals, within and across cortical areas (Fig. 1.E)

Given our model of the microcircuit comprises of exactly two hierarchical areas, “sensory” inputs to the circuit are received at LL4 and outputs are produced at HL2/3. Following the details provided in (38), the relative number of neurons in layers 2/3, 4, 5/6 of an area are in the ratio 5:2:4 respectively. Our base model uses a scaling factor of 16, giving us 80, 32, and 64 neurons in respective layers. All weights at initialization are sampled from 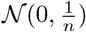, where *n* is the number of neurons.

### 2.2 Tasks

Considering the temporal nature of the world and noting that this structure is key to learning useful representations of stimuli (39), all our tasks are predictive in nature and follow the general structure of learning a sequence 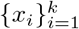. Provided the input elements of the sequence {*x*_1_, *x*_2_, …*x*_*t*_…, *x*_*k* 2_, *x*_*k* −1_} one at a time, the circuit is required to predict the next element at each time step, and cumulatively the sequence {*x*_2_, *x*_3_, …*x*_*t*+1_…, *x*_*k*−1_, *x*_*k*_}. The training objective used is the mean squared error reconstruction loss

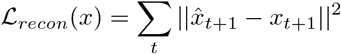

Here 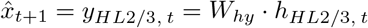 which is the output at HL2/3 at time *t* as given by Eq. 1.

The three tasks in particular that we test on are: i) Sequence memorization (Fig. 1.F), where the microcircuit is trained to learn a given set of sequences, ii) Lattice navigation (Fig. 1.G), where the network needs to predict the next state on a grid given a current state and action, and iii) Binary addition (Fig. 1.H), where the microcircuit is given a binary input and is expected to increment it at every time-step. Full descriptions of all the tasks are provided in Appendix B.

To study the microcircuit in the context of the predictive coding hypothesis and discern if any evidence of error computations can be found explicitly within the circuit, we test our networks on two types of out-of-distribution data (Fig. 1.F) post training: i) Data with temporal violations, where the sequence processed is *x*_1_ − *x*_2_ − *x*_2_ instead of *x*_1_ − *x*_2_− *x*_3_, i.e., the element in the 3^rd^ position is unexpectedly replaced by a repetition of what it saw in the previous time step. ii) Data with additive spatial noise resulting in the input sequence 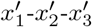 where 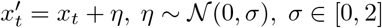.

All our sequences incorporate a repetition of the input stimuli to ensure that every element of the sequence is processed for a sufficient length of time throughout the circuit. As a result, the microcircuit receives the input sequence *x*_1_ − *x*_2_ − *x*_3_ − *x*_4_ as *x*_1_ − *x*_1_ − *x*_2_ − *x*_2_ − *x*_3_ − *x*_3_ and is expected to predict *x*_2_ − *x*_2_ − *x*_3_ − *x*_3_ − *x*_4_ − *x*_4_ as its corresponding output sequence.

### 2.3 Methods for Representational Analyses

The following section gives a brief overview of the methods we use to analyze the geometric properties and informativeness of our representations. Additional explanations and implementation details for all our representational analyses methods are provided in Appendix C.

#### Dimensionality Gain

As per (39), for a given set of representations we define their dimensionality gain (DG) as the ratio of their linear global dimension (L_*dim*_) to their non-linear local dimension (NL_*dim*_), i.e., 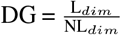. Intuitively, the measure captures whether the representations are encoding simple or complex task-relevant concepts, and if they are doing so in an efficient manner. Moreover, DG of a complex yet structured concept with be high (Fig.2.B - Left), while that of something without intrinsic structure, such as noise, would be low (Fig.2.B - Right).

**Figure 2:**
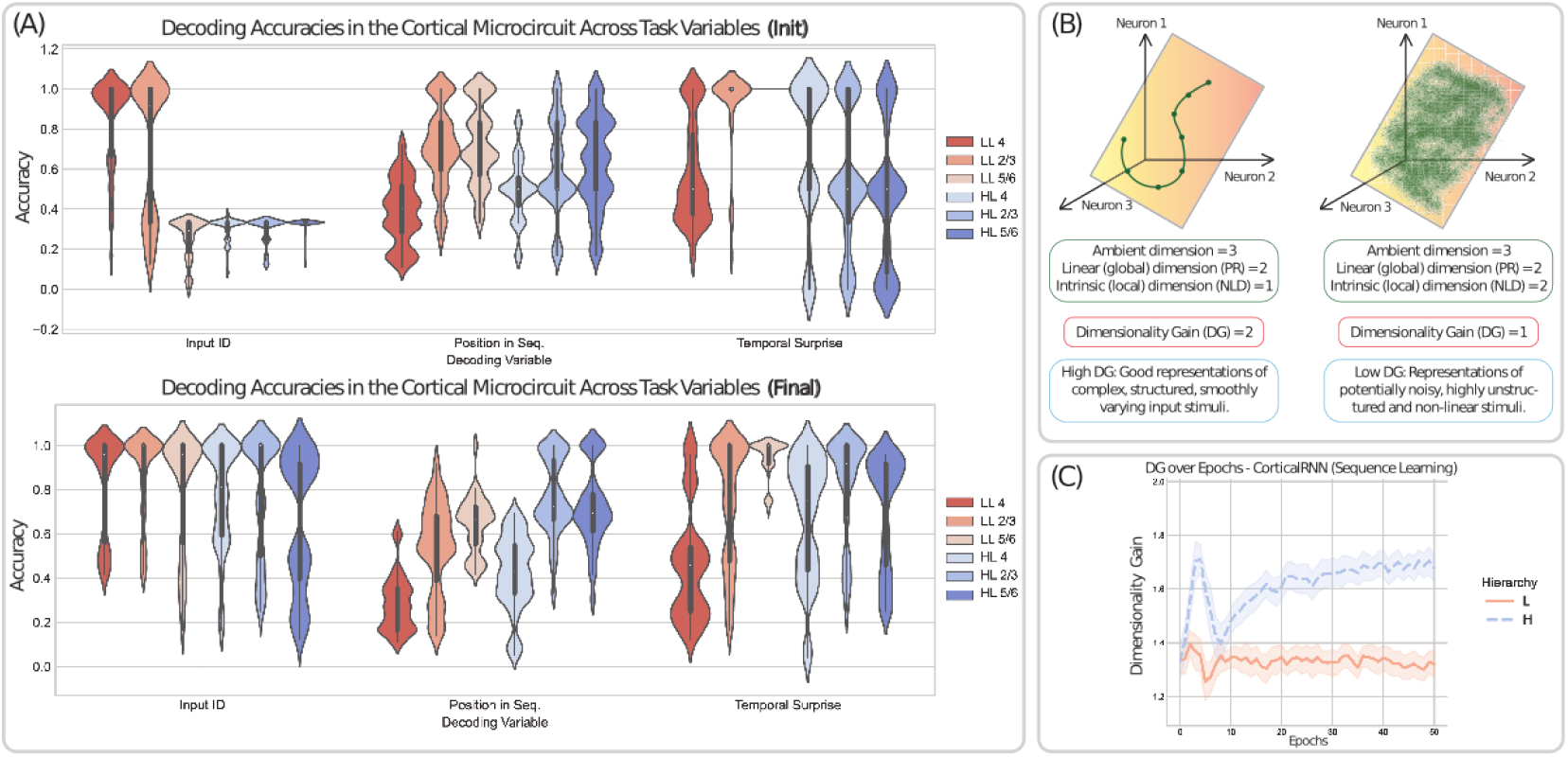
Architectural Biases of the CorticalRNN for the sequence memorization task. (A) Decoding accuracies of task-relevant variables at various layers in the corticalRNN at initialization (top), and after training (bottom). (B) Dimensionality Gain (DG) in the case of a complex yet structured (Left) vs unstructured concept (Right). (C) DG of the corticalRNN during learning of the sequential learning task, split by layers that form the lower cortical area (L - solid line for mean - red) and the higher cortical area (H - dashed line for mean - blue; shaded area implies variance). *(For the purposes of exposition, results in this figure are restricted to the case of the sequence memorization task. Results corresponding to the lattice navigation task and binary addition task are provided in Appendix D*.)

#### Decodability of Task Variables

Another method for measuring the extent to which a set of representations are informative regarding a particular concept is by checking how *predictive* they are of the same, or equivalently, how linearly separable different “classes” relevant to a specific concept are from each other in high-dimensional space. The concepts we test decodability for are: (i) **Input ID** – Which input is being processed by the microcircuit. (E.g., *x*_1_ vs *x*_2_), (ii) **Position** – Where in the sequence you are. (E.g. the first position in the sequence vs the second position in the sequence), and (iii) **Surprise** – Whether the input image being processed is that which is expected or not. (E.g. expected input *x*_2_ presented as a part of a learned sequence vs unexpected input *x*_2_ presented as a repeat that violates a learned sequence)

#### Assessing Neuronal Population Selectivity

Complementing the analyses studying dimensionality of neuronal representations, we also quantify the selectivity of different neuronal populations for a particular concept. To do so, we estimate the minimum number of neurons required to predict a concept from a given set of representations, using the *L*_1_ regularized version of a linear classifier. In doing so, areas that consistently require fewer neurons to predict a concept (i.e., separate classes relevant to that concept) post training with high accuracy can be identified as being more “specialized” or tuned to that particular concept.

## 3 Architectural Biases and Learning in the Canonical Cortical Microcircuit

Given the construction described in Sec. 2.1, in this section we study the effects of the explicit anatomical structure (i.e., inter-areal feedforward and feedback projections) observed in the canonical cortical microcircuit. All results provided are averaged over 5 independent runs of each experiment, performed on a Dell Precision 7920 Tower with an Nvidia Quadro RTX 5000 GPU.

### 3.1 Evidence of Functional Modularization

Observing the decoding accuracies in various layers of the corticalRNN for the sequence learning task, we see that at initialization (Fig. 2.A, top - Left), the identity of the **input** is highly decodable in the lower area in LL4 and LL2/3, while after training (Fig. 2.A, bottom - Left), decodability of the input identity increases to high levels across the microcircuit. In the case of decoding the **position** of the received input vector in the sequence, accuracies are greater in layers higher in the hierarchy, particularly post-training (Fig. 2.A, bottom - Middle). Finally, in the case of decodability of **surprise**, we note that right at initialization, decodability is extremely high for the layers LL2/3 and LL5/6, which both receive feedback from the higher area (Fig. 2.A, top - Right). Post training, accuracies improve to some degree for the layers in the higher areas as well (Fig. 2.A, bottom - Right), however the highest accuracies observed are still at layers LL2/3 and LL5/6 where feedback projections from the higher area directly targeted, overall in agreement with the predictive coding hypothesis. These results collectively make a case for the idea of “functional modularization” within the microcircuit (with similar results for the other two tasks also provided in Appendix D), in that they align with the notion that **lower cortical areas have neuronal representations that correspond to lower-level or “simpler” concepts**, i.e., information that is directly related to the input itself (such as the identity of the input vector or state being received) and **higher cortical areas have neuronal representations that are better tuned to more complex task-related concepts derived from combining different pieces of information corresponding to the task** (such as the temporal position of an input within a sequence). These results are further supported by the dimensionality gains (Fig. 2B) that we see averaged over the populations in the lower and higher processing areas separately (Fig. 2C). Over the course of learning, we note that the average DG of the lower area remains relatively constant, implying that it encodes simpler information that has a constant ratio of ambient and latent dimensionality in neuronal space. On the other hand, that of the higher area increases with learning, implying that it encodes information that is higher dimensional in the ambient sense while still being structured and encodable with few latent variables.

### 3.2 Inductive Biases due to Inter-Areal Feedback and Time Delays in the Cortical Microcircuit at Initialization

Further delving into the results we obtain for decoding task-related concepts, we observe that areas receiving inter-areal feedback, viz. (*LL*2*/*3, *LL*5*/*6) are particularly adept at decoding surprise or unexpectedness in the sequence memorization task. Moreover, they do so with high accuracy not only post training, but at initialization as well (Fig.3A (Left), Fig.3B). We posit that this result is a consequence of the presence of time-delays in the projection of inter-areal communication and the easily separable nature of our stimuli. More concretely, we hypothesize and subsequently show that the microcircuit exploits the sequential nature of the task and separability of the input as follows:

**Figure 3:**
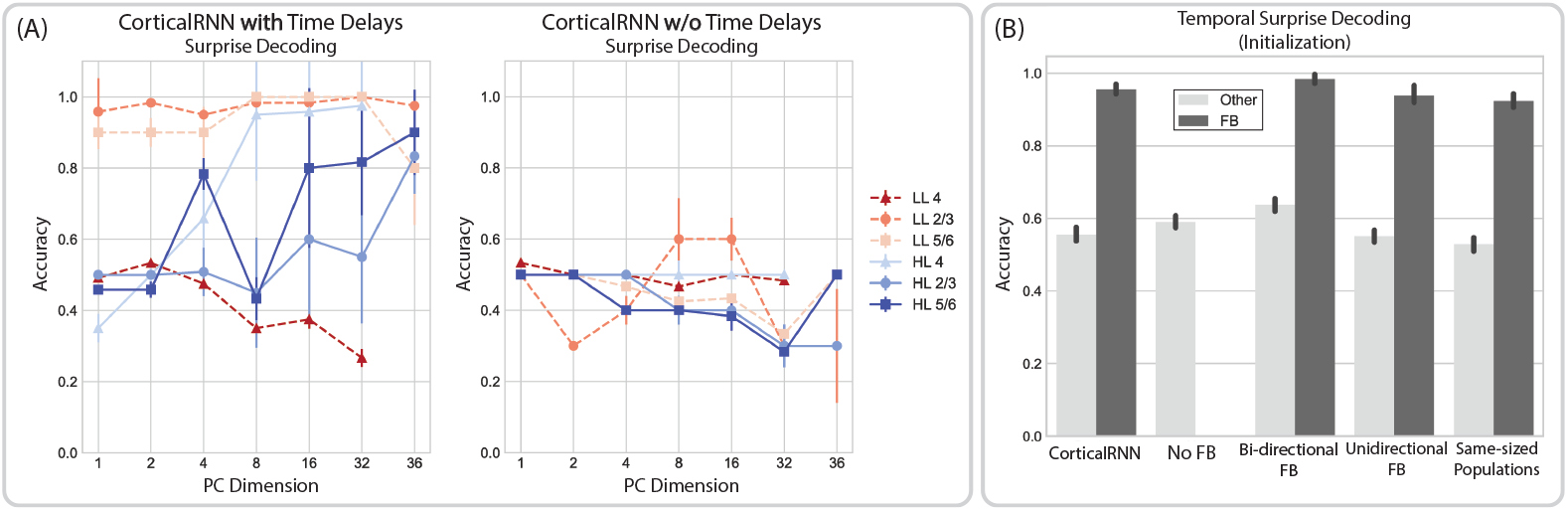
Temporal surprise decoding in the CorticalRNN for sequence memorization. (A) Decoding of temporal surprise at different layers of the corticalRNN with (Left) and without (Right) time delays, using the principal components of their representations at initialization. (LL4, HL4 have 32 neurons.) (B) Effects of inter-areal time delays on decodability of surprise at initialization, in layers receiving inter-areal feedback (dark grey) vs not (light grey), in the CorticalRNN and its structural variants.

1. The first three elements of our expected sequences are always permutations of the inputs *x*_1_, *x*_2_, *x*_3_, which ensures that all inputs are seen as both expected and unexpected in the same temporal position as part of some sequence. E.g., both *x*_1_−*x*_2_−*x*_3_−*x*_4_ and *x*_1_−*x*_3_−*x*_2_−*x*_5_ belong to a set of expected sequences. The standard sequence *x*_1_ − *x*_2_ − *x*_3*e*_ − *x*_4_ has the input *x*_3_ as expected in the third position, but the sequence *x*_1_ − *x*_3_ − *x*_3*v*_ − *x*_5_ is temporally violated from its natural form *x*_1_ − *x*_3_ − *x*_2_− *x*_5_ and sees the input *x*_3*v*_ unexpectedly, again in the third position. (Note that *x*_3_ = *x*_3*e*_ = *x*_3*v*_ in input space; we make the distinction between them simply for the purposes of exposition.)
2. Decoding surprise in the sequence memorization task requires the ability to find a separating hyperplane between inputs that are received as expected in a sequence vs. those that are received unexpectedly, i.e., being able to distinguish *x*_3*e*_ from *x*_3*v*_, in neuronal space with a separating hyperplane.
3. When a time delay is present in inter-areal projections, the representations formed in populations which receive feedback are a combination of the inputs being received at that particular timestep, as well as that received from 2 timesteps prior. However in the absence of any inter-areal time delays, the entire microcircuit simply processes the input being received at that particular timestep.
4. Given that we provide repeated presentations of the inputs to the microcircuit to represent their duration, i.e., our input sequences are *x*_1_ − *x*_1_ − *x*_2_ − *x*_2_ − *x*_3*e*_ − *x*_3*e*_ and *x*_1_ − *x*_1_ − *x*_3_ − *x*_3_ − *x*_3*v*_ − *x*_3*v*_, in the presence of time delays the neuronal populations *LL*2*/*3, *LL*5*/*6 process a combination of the inputs *x*_2_,*x*_3*e*_ in the case of the standard sequence and *x*_3_, *x*_3*v*_ in the case of the violated sequence at the third temporal position. In the absence of time delays, at the third temporal position, the microcircuit simply processes *x*_3*e*_ in the standard case and *x*_3*v*_ in the temporally violated case at all layers.
5. Noting that all the weights of the microcircuit are Gaussian-distributed at initialization and that there are no projections that lead to any drastic compression or expansion across layers or areas (i.e., by an order of magnitude or greater), the relative distances between inputs in the input space are largely preserved in the representational space, following the Johnson-Lindenstrauss lemma (40; 41).
6. Assuming that the inputs are indeed typically fairly separable in input space, this explains why populations which receive inter-areal feedback can indeed distinguish an expected input from an unexpected one in the presence of time delays; these layers have representations that easily allow a separating hyperplane between representations of the form ≈ *x*_2_ + *x*_3_ for the expected *x*_3*e*_ vs those of the form ≈ *x*_3_ + *x*_3_ for the unexpected *x*_3*v*_. However in the absence of time delays, this would not be the case as the representations to be separated would both correspond to the same input ≈ *x*_3_ for both the expected and unexpected *x*_3_.

More formal, mathematically-grounded justifications supporting our hypothesis and subsequent rationale are provided in Appendix E. In particular we show that in the case of the sequence memorization task with input Bernoulli vectors ∼ ℬ (*p*), our inputs are indeed separable. Furthermore, given the structure of the cortical microcircuit and assuming all weights are initialized from 𝒩 (0, 1), this separability can be maintained in representation space within a multiplicative factor of distortion *ϵ*, provided that the minimum dimension that our inputs are be projected to is 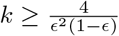 ln 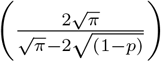 .

Our empirical results (Fig.3) validate our intuitions and theoretical justifications, where we see that it is indeed the structure of the stimuli in input space, sequential nature of the memorization task, and intrinsic time delays in the microcircuit that explain our results (Fig.3.A). Furthermore, the ability to distinguish temporal violations with high accuracy is specific to layers which receive inter-areal feedback across different types of architectures (Fig.3.B), making a strong case in favour of the predictive coding hypothesis. While we make the above argument in the case where our stimuli are highly separable in input space, the idea truly is more general and would hold in whichever cortical area the representations corresponding to a stimulus are distinguishable. Hence, **these results suggest a general motif of cortical computation underlying the efficient information processing in sequential tasks** that is often observed across various natural environments and animals (42).

Finally, we note that architectures receiving feedback are slightly more robust than the no-feedback architecture to temporal violations post-training (Appendix F), i.e., when given a temporally violated input sequence, they still produce outputs that are correspond to the original sequence more consistently. However, in the presence of spatial noise, we find no significant differences in the performance across the various architectures across tasks.

## 4 A Predictive-Coding Inspired Training Strategy

While the previous section focused on the effects of structural priors (i.e., anatomical constraints) on how information is processed within the canonical microcircuit, we now study the effects of imposing a *functional prior* while learning. In particular, we look to test whether the predictive coding hypothesis when incorporated in conjunction with the structure of the microcircuit leads to evidence supporting the notion that areas receiving inter-areal feedback are the seats of error computation. Our training strategy inspired by predictive coding (1) involves two distinct phases, each updating the model with different loss functions. In the first phase, like before, the entire network is trained using a reconstruction-based loss of the form

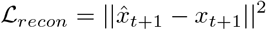

In the second phase, only parts of the network responsible for generating and transmitting prediction signals are trained. These are the higher cortical layers *HL*2*/*3, *HL*4, *HL*5*/*6, and feedback connections 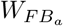 and 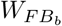. The training now uses a predictive-coding (PC) loss, calculated as the average of the prediction errors from the two feedback projections *FB*_*a*_ and *FB*_*b*_ as defined in Section 2.1. Each prediction error is calculated as the difference between the signal conveyed through the feedback connection and the neuronal representation of the layer to which the signal is directed. Specifically, we minimize

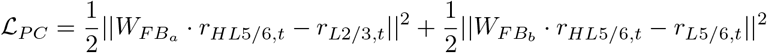

We then alternate between the two phases, so that parameters in the higher areas are encouraged to be predictive of the activity in the lower area. The explicit training algorithm is provided in Appendix G.

### 4.1 Effects of Predictive-Coding Inspired Training on Representational Dimensionality Across Layers and Areas

Studying DG trends across training in various layers of the microcircuit while using only a reconstruction-based loss shows no appreciable differences across various architectures (Fig. 4.A). The DGs of layers in the higher processing area are greater than those in the lower area, and this trend is maintained across all architectures. Moreover, addition of the PC loss does not affect the general trend of DGs across areas for architectures that have no feedback connection from the higher area to LL5/6 either (Fig. 4.B, Top - (ii), (iii)). However, addition of the PC loss significantly changes the DG of LL5/6 in architectures where the layer does receive feedback (Fig. 4.B, Top - (i), (iv), (v)). In particular, we note that in these cases the DG of LL5/6 drops with training, implying that the representations in the layer in these cases has linear and non-linear dimensionalities that grow in tandem. This consequently points to the idea that the phenomenon being encoded in LL5/6 lacks smooth structure and predictability. We therefore hypothesize that **LL5/6 in these cases encodes “surprise” as posited by predictive coding theory, which by definition is unstructured and unpredictable. Furthermore, this encoding of surprise in the area is facilitated by both, the physical feedback connection as well as the training objective**, and therefore remains conspicuously absent in the dimensionalities of representations in LL5/6 of the architecture without any feedback connections (Fig. 4.B, Top (ii) and (iii)). While Fig. 4 shows these trends in the sequence learning task, we find that these results hold consistently across other tasks as well (Appendix H).

**Figure 4:**
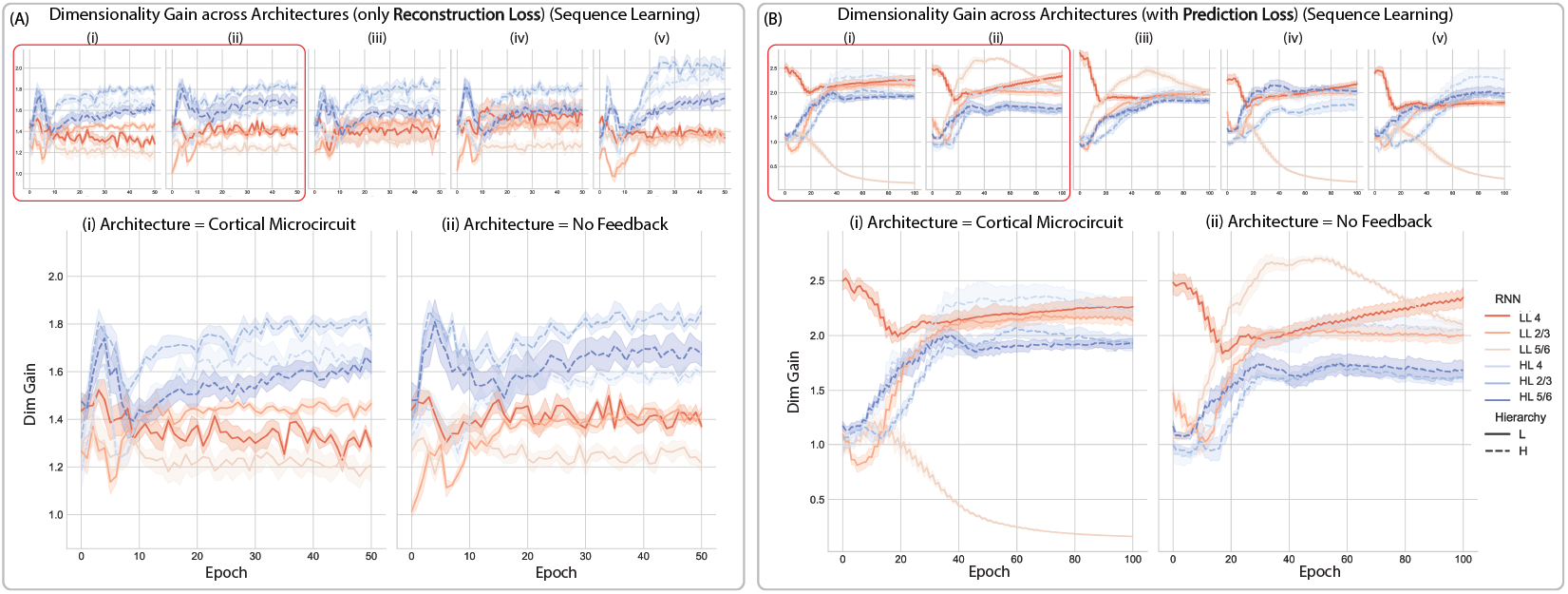
Effects of Using Predictive-Coding Based Training Objective on Dimensionality Gain. Panels (A) and (B) are juxtaposed to show the differences that arise in the DG of various layers across learning (A) without and (B) with the inclusion of a predictive-coding based objective respectively. In both, the top row shows DG across layers for architectural variations in the following order: i) canonical cortical microcircuit, ii) no feedback, iii) bi-directional feedback, iv) uni-directional feedback, v) population-controlled microcircuit. The lower panels zoom into the DGs of the (i) cortical microcircuit and (ii) no-feedback architectures.

### 4.2 Effects of Predictive-Coding Inspired Training on Neuronal Specialization

Following our observations of how the DG changes in LL5/6 when trained with the PC loss, we looked to verify our hypothesis that this was indeed caused due to the neurons in the population encoding surprise. To do so, we checked the neuronal selectivity for temporal surprise in the neurons. By requiring a sparse set of neurons to effectively identify surprise elements vs those which are expected implies that the concept of surprise is represented efficiently in that population of neurons, and is what largely drives their activity. We find that our results support our hypothesis, and that the **addition of the PC loss results in fewer neurons being required to encode surprise in LL5/6** (Fig. 5B - Right) as compared to when not using the loss (Fig. 5A - Right), but approximately the same number of neurons are needed to do so at initialization in both cases (Left - Fig. 5A & B). These results hold over multiple runs and are provided in (Appendix I).

**Figure 5:**
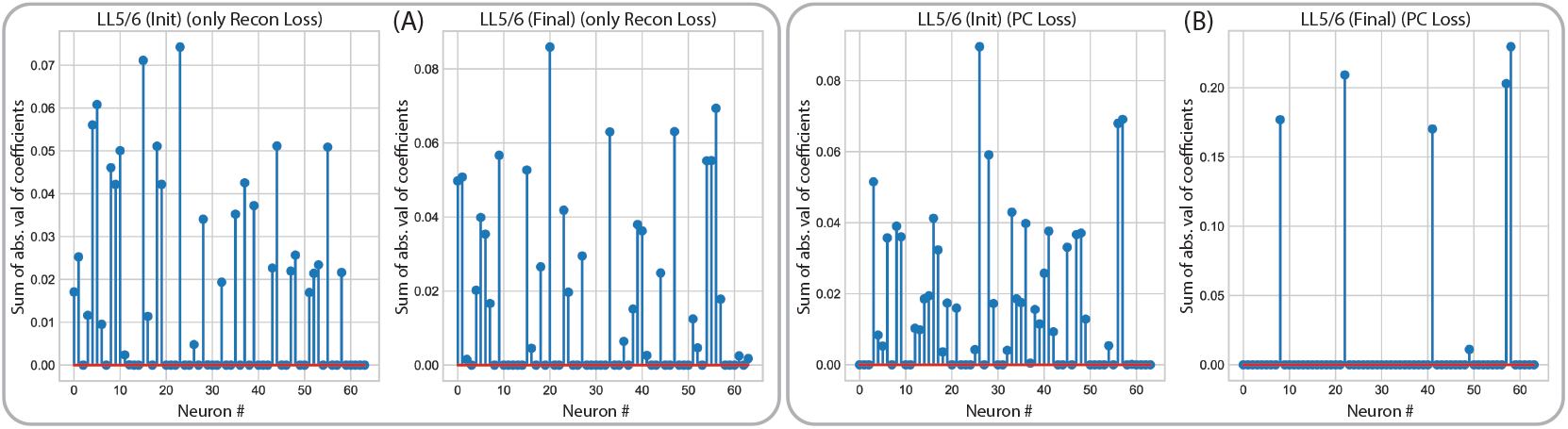
Effects of Using Predictive-Coding Based Training Objective on Neuronal Selectivity. Examples of neurons used to decode surprise in LL5/6 (A) without, and (B) with the PC loss.

## 5 Conclusion

Our work provides compelling evidence that indeed the architectural constraints of the canonical cortical microcircuit influence information representation within it, and also that functional priors in terms of how the network is trained play a key role in its learning and the geometry of its neuronal representations. Specifically, we find that i) the structural primitives (i.e., feedback projections) of the network and the nature of sequential tasks can strongly influence which areas best represent certain task-relevant concepts, and ii) both physical structure and training objective can independently influence the dimensionalities of representations across the microcircuit.

That said, there remain a number of open questions and avenues for further interrogation; For one, our setting is rather limited in that we study the microcircuit with only two hierarchically-related areas. How results might change with increasing “depth” is unknown. Likewise, the effects of incorporating additional biological elements into the model, e.g., different cell-types, Dale’s law, as well as local credit assignment schemes might be significant. Finally, we also note that our work assumed the canonical microcircuit structure and studied learning in it subsequently – but would it be possible to identify physiological phenomena that might constrain development in the cortex and reverse-engineer how this structure might emerge organically? Answers to these questions would go a long way in not only understanding the brain, but could also pave the way for more efficient, robust, and interpretable ML/AI systems by leveraging the appropriate architectural and functional priors.

## Code Availability

Code for models and analyses included in the manuscript is available at: https://hchoilab.github.io/corticalRNN.

## Acknowledgments and Disclosure of Funding

This work was supported by the National Eye Institute of the National Institutes of Health under Award Number R00 EY030840. The content is solely the responsibility of the authors and does not necessarily represent the official views of the National Institutes of Health.

## A Layer-wise computations at an arbitrary time step

The input to the microcircuit at an arbitrary timestep *t* is given by *x*^(*t*)^. Recurrent weights at layer *a* are written as *W*_*a*_ and its neural activity at time *t* is written as 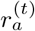.

Weights of the intra-areal backbone (i.e., weights connecting layers in the same cortical area) are given as *W*_*BB*_. On the other hand, *W*_*FF*_ represents weights corresponding to the feedforward projection from the lower cortical area to the higher cortical area. *W*_*FBa*_ and *W*_*FBb*_ represent weights responsible for the feedback projections from the higher cortical area (HL5/6) to the lower cortical area, to LL2/3 and LL5/6 respectively.

Weights projecting the sensory input *x*_*t*_ to the recurrent microcircuit representing the lower cortical area are given by *W*_*ih*_. Projections from layer *a* to layer *b* are denoted as *proj*(*a* → *b*).

The *tanh*(*·*) and ReLU non-linearities are denoted by *ϕ*(*·*) and (*·*)_+_ respectively.

Computations carried out at various layers at any given timestep *t* are subsequently given as follows:

### Lower Layer 4 (LL4)

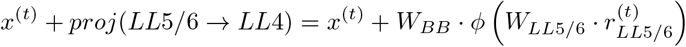

### Lower Layer 2/3 (LL2/3)

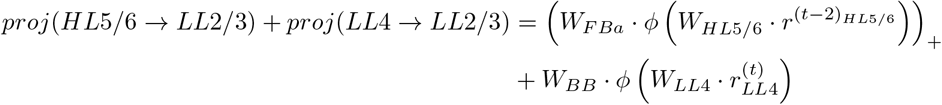

### Lower Layer 5/6 (LL5/6)

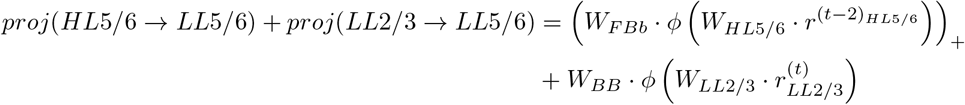

### Higher Layer 4 (HL4)

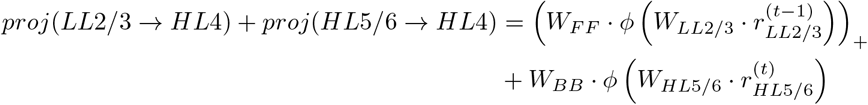

### Higher Layer 2/3 (HL2/3)

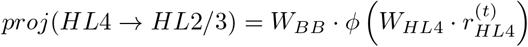

### Higher Layer 5/6 (HL5/6)

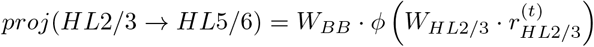

## B Task details

### Sequence Memorization

Our exemplary task (Fig. 1.F) entails memorizing a set 𝒮 with elements that are sequences of length *k*. In particular, 𝒮 is defined as the collection of all (*k* − 1)! possible sequences where the first *k* − 1 elements of the sequence are permutations of *{x*_1_, *x*_2_, …*x*_*t*_, …*x*_*k*−1_*}*, and each *x*_*t*_ is a sparse vector such that *x*_*t*_ ∈ *{*0, 1*}*_*D*_ where 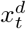 ∼ Bernoulli(*p*), *d* = *{*1, …*D}* and *t* = *{*1, …*k}*. The *k*^*th*^ element is a sequence identifier or label drawn from the same distribution as the other 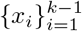. E.g., when *k* = 4, *𝒮* = *{*(*x*_1_, *x*_2_, *x*_3_, *x*_4_), (*x*_1_, *x*_3_, *x*_2_, *x*_5_), (*x*_2_, *x*_1_, *x*_3_, *x*_6_)…, (*x*_3_, *x*_2_, *x*_1_, *x*_9_)*}* and |*𝒮*| = 6. The inputs are sampled such that 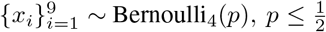. The training and test sets subsequently consist of sequences 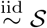. We use *n*_*train*_= *n*_*test*_ = 1000. With *p* = 0.25, *k* = 4, and *D* = 64, the sets of expected and unexpected sequences take the following form:

**Table 1:**
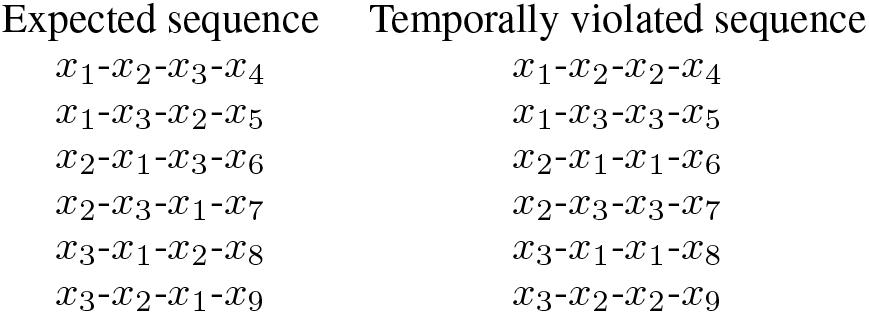
Expected and unexpected sequences for the memorization task.

#### Lattice Navigation

Our second task is similar to that of (39), requiring the network to predict the next state *x*_*t*+1_ it would traverse to on an *n × n* grid, given a sequence of the previous states (*x*_0_, *x*_1_, …*x*_*t*_) along with the actions (*a*_0_, *a*_1_, …*a*_*t*_) taken at each of those states (Fig. 1.G). Each position on the grid corresponds to a fixed high-dimensional state *x*_*t*_ ∈ *{*0, 1*}*_*D*_ and the actions *a*_*t*_ ∈ 1_*i*_, *i* = *{*1, 2, 3, 4*}* are one-hot vectors corresponding to the four cardinal directions.

In the case of temporal violation at a timestep *t*, we provide the microcircuit with the appropriate action, but repeat the state vector *x*_*t*−1_ as the state input *x*_*t*_.

For our purposes, *D* = 64, *n*_*train*_ = 1600, *n*_*test*_ = 400, *n*_*epochs*_ = 50. *p* = 0.75

#### Binary Addition

Our last task requires the microcircuit to interpret a binary input in *D* bits and increment it by one at every time step, also in binary (Fig. 1.H). Specifically, given the binary representations (*x*_*z*_, *x*_*z*+1_, *x*_*z*+2_, …) of the numbers (*z, z* + 1, *z* + 2, …) where *x*_*t*_ ∈ {0, 1}^*D*^ and *z* ∈ ℝ, the expected output is (*x*_*z*+1_, *x*_*z*+2_, *x*_*z*+3_, …) i.e., the corresponding *D* bit binary representations of the numbers (*z* + 1, *z* + 2, *z* + 3) at each corresponding time step.

In the case of temporal violation at time-step *t*, we repeat the input vector *x*_*t* − 1_ instead of presenting the microcircuit with the appropriate input *x*_*t*_.

For our purposes, *D* = 16, *n*_*train*_ = 4000, *n*_*test*_ = 1000, *n*_*epochs*_ = 50.

For training on all our tasks we use the Adam optimizer (43) with a step size of 0.001, in its standard PyTorch (44) implementation.

## C Representational analyses details

### Dimensionality Gain

DG captures whether representations encode simple or complex task-relevant concepts, and if they are do so in an efficient manner. This can be reasoned about as follows:

- If a complex concept is being encoded by a set of neurons, their linear or *global* dimension is high as the data occupy much of the neuronal space available to them. However, since the information in the representations is relevant to the task, there is structure reflected in them that is captured by a non-linear transformation, therefore resulting in a low non-linear or *local* dimension. As a result, the DG of such a concept is high (Fig.2.B - Left).
- On the other hand, if a concept is simple, both its linear and non-linear dimesionalities are low, making the DG low as well.
- Finally, if the information encoded in the representations has no real structure, the linear and non-linear dimensionalities both tend to be high as the representations seemingly occupy all of the ambient space, again making the DG low (Fig.2.B - Right).

As a result, we can think of increasing DGs to be a sign of learning, with higher values corresponding to more complex phenomena, while lower or decreasing DGs indicate lack of structure or noise being encoded in the representations.

Our measure of linear dimenisonality (L_*dim*_) is the participation ratio (PR) (45; 46) where

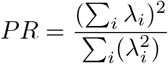

and *λ*_*i*_ are the eigenvalues of the covariance matrix of the neuronal representations.

Non-linear measures of dimensionality however are far less standardized, and therefore we take the average of four different measures, viz., CorrDim, MLE, DANCo, and MiND_ML_ (47; 48; 49; 50) as our measure for non-linear dimension (NL_*dim*_) from the `scikit-dimension` python package (51).

Consequenlty,

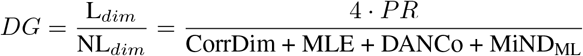

### Linear decoding of task variables

A greater degree of linear separability amongst classes of a concept when using fewer dimensions implies the representations are highly tuned to the particular concept, and therefore more informative of the concept under consideration. We therefore compute the principal components (PCs) of the neuronal representations from the different layers, and subsequently fit separating hyperplanes between the PCs corresponding to different classes pertinent to the various concepts. The separating hyperplanes are found using a standard linear support vector machine, without any modifications from the `scikit-learn` python package (52).

### Neuronal population selectivity

To quantify how tuned the neuronal activity in different cortical layers are to specific concepts, we find the most sparse subset within a given population that allows us to draw separating hyperplanes between different classes relevant to the concept (e.g., different input identities in the case of decoding input states, and expected vs unexpected inputs when decoding surprise) as accurately as the full population itself. The rationale behind doing so is that doing so allows us to find the neurons which dominate the activity relevant to a particular concept, while also accounting for redundancies within the activities of the neurons themselves. Assuming a sparse multinomial logistic regression model trained with cross-entropy, our objective is

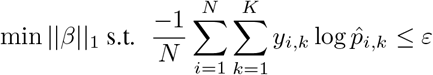

where *ε* is the error of the classifier when using all neurons. *y* is the true label, *K* is the number of classes relevant to the concept being tested, and *N* is the total number of samples. As with typical logistic regression, 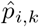 is the predicted probability that the *i*^*th*^ sample belongs to class *k*, given by

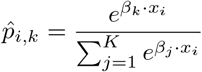

*β* ∈ ℝ^*d*^ is the coefficient vector and *x* ∈ ℝ^*d*^ the firing rates of the *d* neurons in a neuronal population.

## D Decoding task-relevant variables

### D.1 Binary prediction task

**Figure 6:**
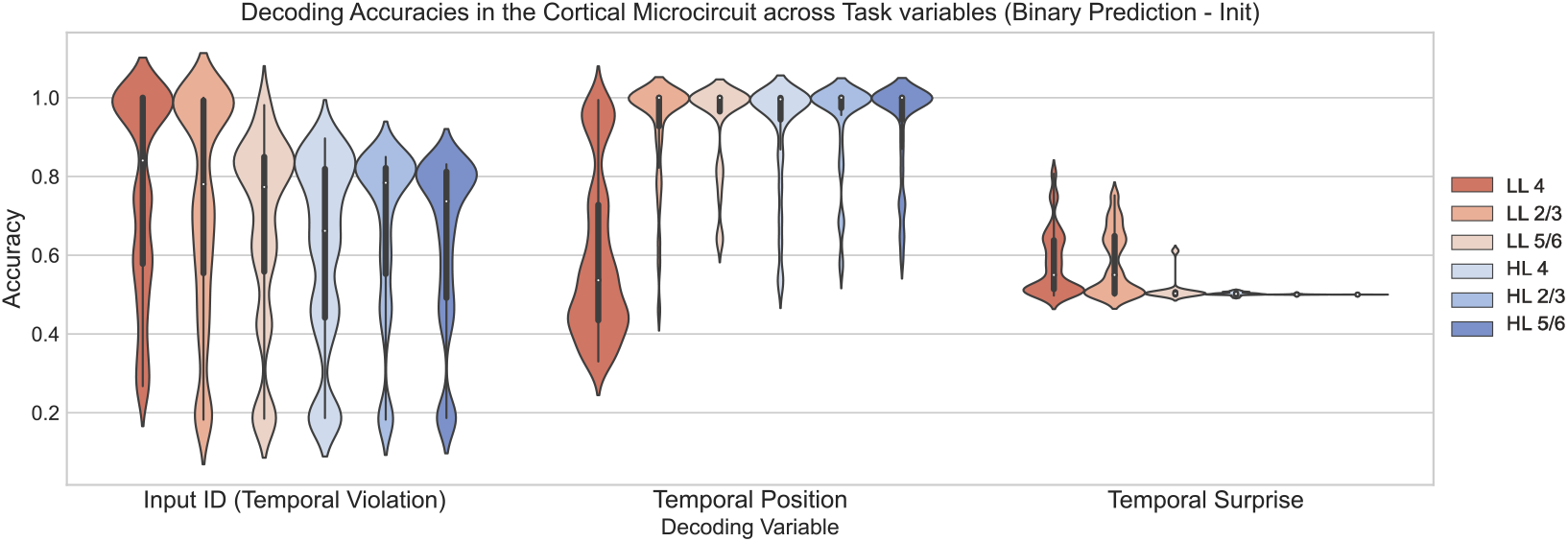
Binary prediction (Initialization): Decoding of task-relevant variables

**Figure 7:**
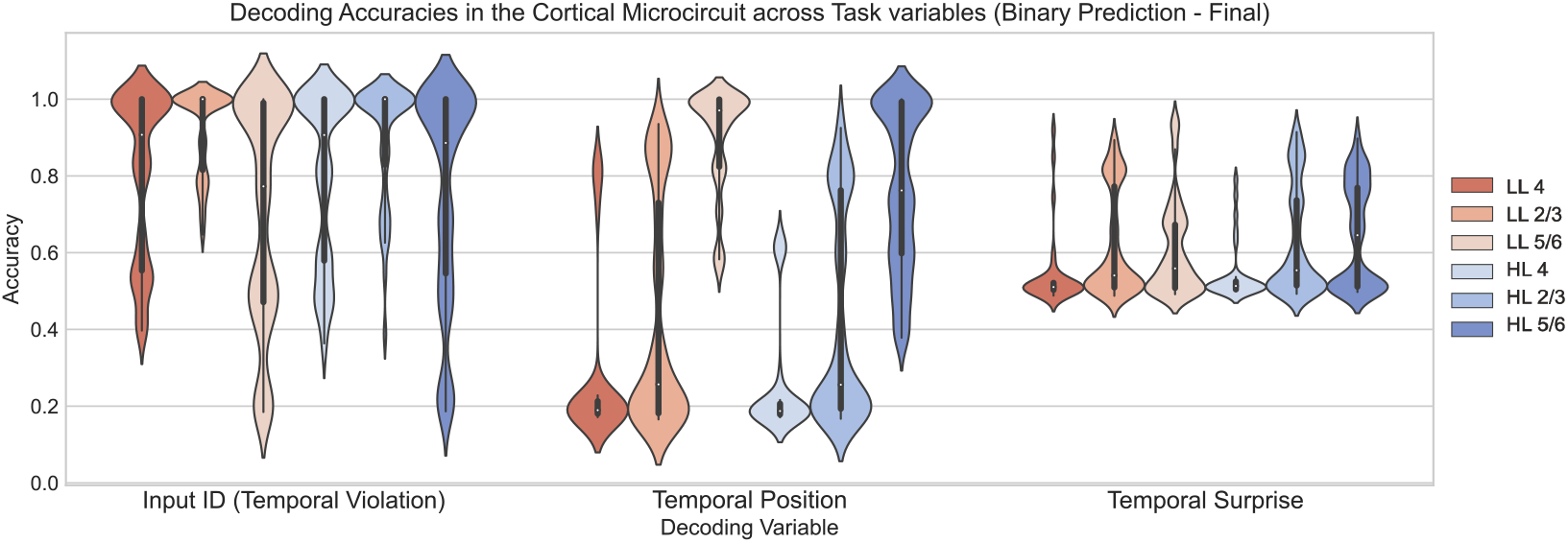
Binary prediction (Final): Decoding of task-relevant variables

As before in Sec. 3.1, we notice that input identity is more decodable in layers in the lower area at initialization (Top - Left), and remains so even after training (Bottom - Left). At the same time, the input ID becomes highly decodable in the higher area too with training (Bottom - Left). On the other hand, position of an input in the sequence is most decodable in layers higher in the hieararchy, along with those receiving feedback at initialization (Top - Middle), while post training LL5/6 and HL5/6 seem to do best (Bottom - Middle). At initialization all layers seem to only decode surprise at chance (Top - Right), but post training accuracies improve across the microcircuit, with LL5/6 reaching the highest accuracies (Bottom - Right). On the whole, the trend that simpler concepts are better encoded in the lower area while more complex ones are better encoded in layers belonging to the higher cortical area still holds.

### D.2 Lattice navigation task

**Figure 8:**
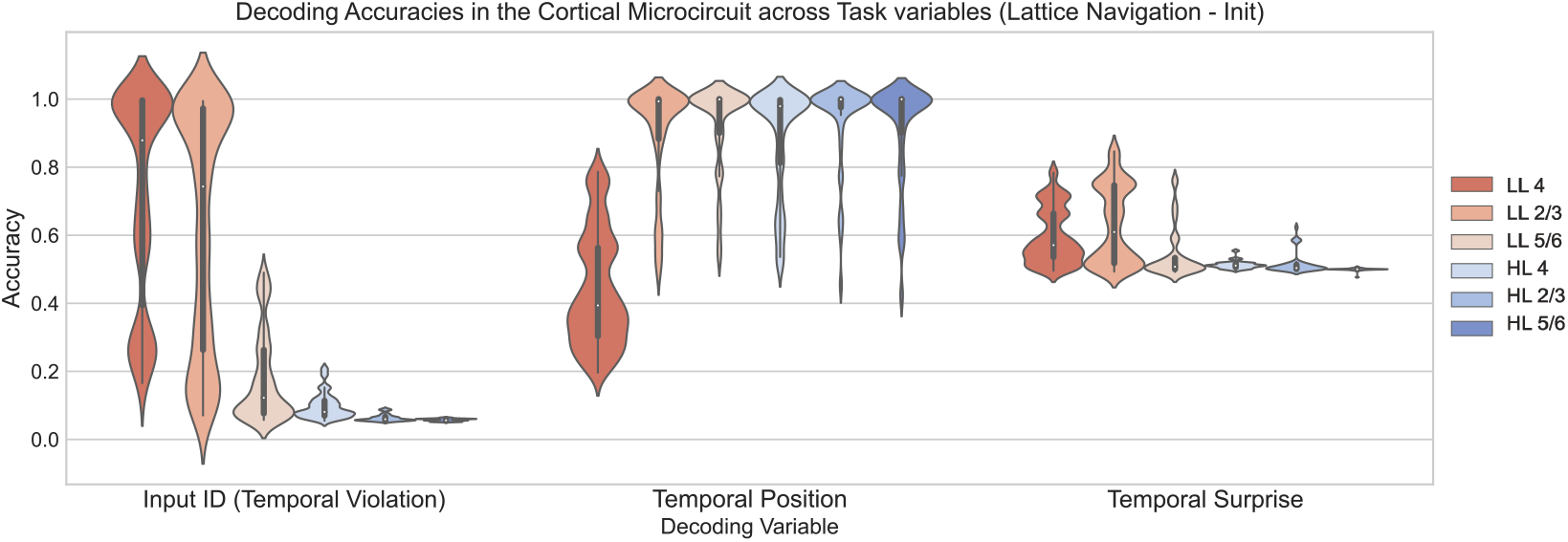
Lattice navigation (Initialization): Decoding of task-relevant variables

**Figure 9:**
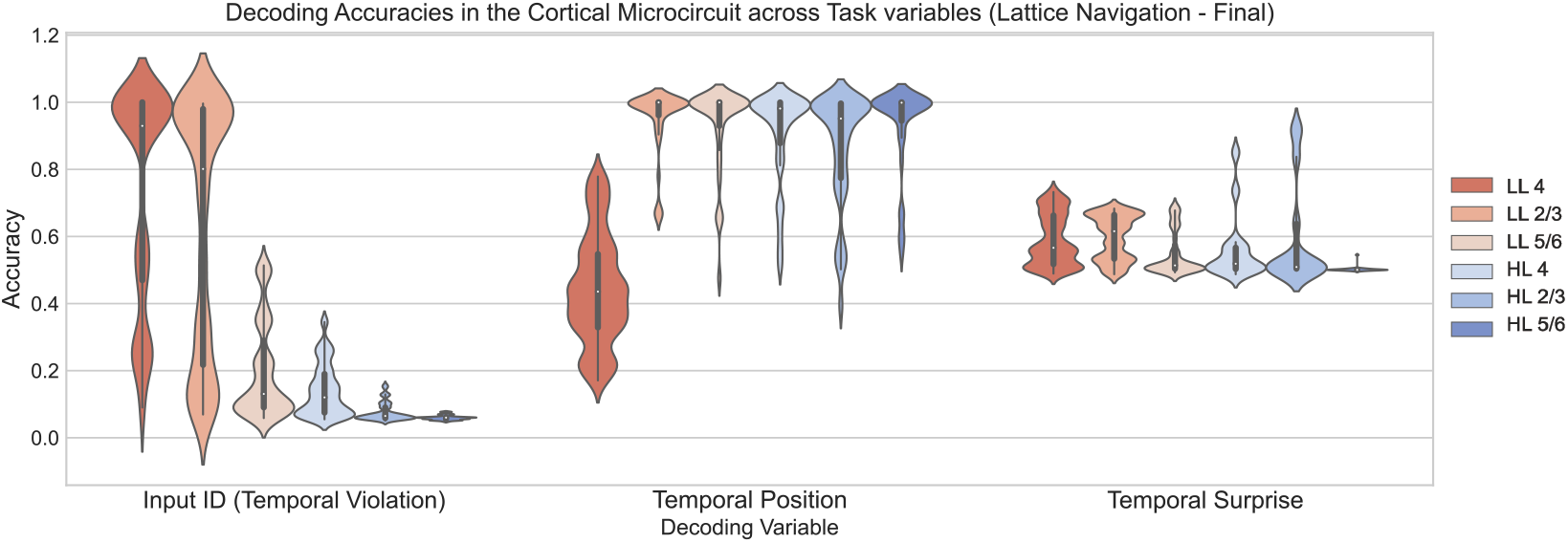
Lattice navigation (Final): Decoding of task-relevant variables

Functional modularization is extremely evident in the case of input image decoding (Left - top & botton) and position decoding (Middle - Top & Bottom). Surprise decoding at initialization (Top - Right) was best performed by LL2/3, but post-training overall decodability for surprise across the microcircuit seems to be reduced (Bottom - Left).

## E Surprise decoding: Effects of feedback and inter-areal time-delay at initialization

In this section we provide mathematically-grounded justifications for the intuitive explanations provided in Sec.3.2 regarding the decodability of surprise at initialization in *LL*2*/*3, *LL*5*/*6 for the sequence memorization task. Our analysis also explains why there is no significant separability in the representations of these layers in the absence of any such time-delays. In particular, we show more rigorously that

1. Our input stimuli are separable in input space, and this separability is maintained in the case of expected sequence memorization but not in the case of temporal violations.
2. The probability of a random vector being able to act as a linear separator (i.e., decoder) increases with vectors that are farther apart and therefore easily separable.
3. The separability amongst stimuli in the input space largely carries over to neuronal space (specifically in *LL*2*/*3, *LL*5*/*6) at initialization.

### E.1 Separability of Bernoulli vectors in input space

Given *x, y* ∼ ℬ (*p*) where *p* is the probability that any element *x*_*i*_ or *y*_*i*_ = 1, and *x, y*∈ {0, 1}^*d*^ with *d* being the length of the vector, the expected distance between the two vectors *x, y* is:

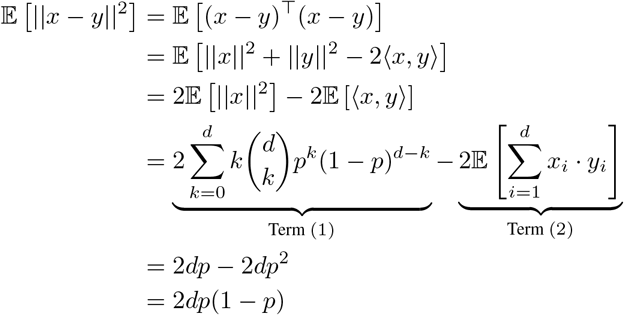

Term (1) is obtained by first making the substitution 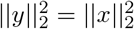 as their expected values are the same given that they are both drawn from the same distribution, and then summing them. Next, we note that the expected norm squared of the vector *x* is the exact same quantity as the number of ones in the arbitrary vector *x*. This is the same quantity as the expected value of successes in a Binomial random variable with parameters (*d, p*), which in turn is equivalent to the sum of *d* independent Bernoulli trials *X*_*i*_ with the parameter (i.e., mean) *p*, resulting in the following simplifications

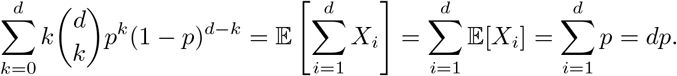

Term (1) consequently can be written as 2*dp*.

Term (2) on the other hand is the expected value of the sum of the element-wise multiplication of the vectors *x, y*. This is the same as counting the number of matching elements between the two vectors, while noting that every element is drawn independently from ℬ(*p*). Therefore, we have

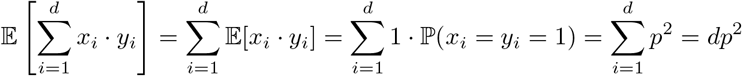

which simplifies Term (2) to 2*dp*^2^.

In the sequence memorization task, substituting *x* = *x*_2_ + *x*_3_ and *y* = *x*_3_ + *x*_3_ for the case of the expected sequences as discussed in 3.2, we see that the same squared distance 2*dp*(1−*p*) is maintained by any two arbitrary vectors *x*_2_, *x*_3_ in input space^5^ as 𝔼 [ ||*x*− *y*|| ^2^] = 𝔼 [||(*x*_2_ + *x*_3_) − (*x*_3_ + *x*_3_) ||^2^] = 𝔼 [ ||*x*_2_ − *x*_3_ ||^2^] and their eventual representations in neuronal space at *LL*2*/*3, *LL*5*/*6, assuming that the same distance is largely preserved in representational space.

But before showing the latter, we will first confirm that an increased distance between vectors implies a higher probability of being able to draw a separating hyperplane between them.

### E.2 Probability of finding a separating hyperplane between *δ*−vectors

Without loss of generality, assume vectors *x, y* ∈ ℝ^*d*^ are both unit-norm and the distance between them is *δ*. The probability then, that a random vector *v* can separate the two is

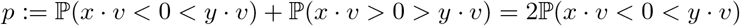

Assuming uniformly sampled *v* ∈ *𝕊*^*d*−1^ we have *v* is equal in distribution to the vector 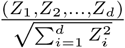 where *Z*_*i*_ are are independent standard normal random variables. Additionally, we note that the dot product of a Bernoulli vector ∼ ℬ (*p*_*b*_) with a random vector *v* sampled from the unit sphere follows the same distribution as *v*, as the dot product simply samples a subset of the vector *v*. Consequently,

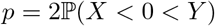

where 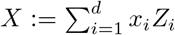 and 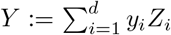. The random variables *X* and *Y* are jointly normal with zero means, variances 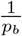, and correlation *r* such that

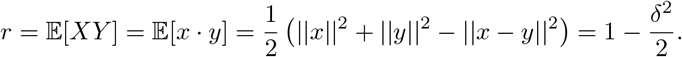

The pair (*X, Y* ) equals 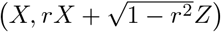 in distribution, where *Z* is a standard normal random variable independent of *X*. So,

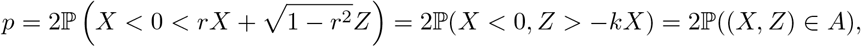

where 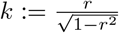 and *A* is the angle between *{*(*x*, 0) : *x* ≤ 0*}* and *{*(*x*, −*kx*) : *x* ≤ 0*}*.

Since the distribution of the random vector (*X, Z*) in ℝ^2^ is rotation invariant, we conclude that the probability in question is 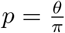, where the measure of the angle *A* is

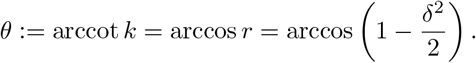

In particular, it follows that 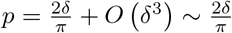 as *δ <<* 1.

### E.3 Johnson-Lindenstrauss lemma

To make our final point that the distances between vectors in input space are largely preserved in representational space at initialization, we invoke a modified version of the Johnson-Lindenstrauss lemma as stated by **Theorem 2 (Neuronal RP)** in (41).

Let *x, y* ∈ *ℝ*^*d*^ and let *x*^*′*^, *y*^*′*^ be their projections onto ℝ^*k*^ via a random matrix *R* whose entries are chosen independently from 𝒩 (0, 1). Then,

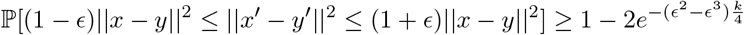

The above statement guarantees that a standard normal random Gaussian projection can preserve pairwise distances ||*x* − *y*|| ^2^ upto an arbitrarily small precision *ϵ* with probability 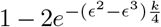 as long as we project onto a minimum dimension *k*. We therefore can use the above result to guarantee that inputs that are appreciably far apart in input space will maintain similar distances (and hence separability) in representational space, given that they’re not projected onto a dimension that is significantly smaller than what *k* should be.

We note however that there are two inter-areal projections which occur to the input before it is processed by either *LL*2*/*3 or *LL*5*/*6 as part of the feedback signal, since it is first projected forward to *HL*4 and then projected back from *HL*5*/*6. Therefore, allowing for the representation to change by a multiplicative factor of *ϵ* at each step, we have

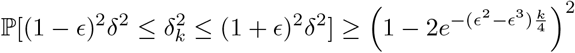

where *δ* = ||*x* − *y*|| is the original distance between the two vectors *x, y* and *δ*_*k*_ is the distance between the projected vectors *x*^*′*^, *y*^*′*^ in *k*−dimensional space. If we let 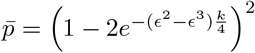 we find that the minimum dimension *k* which we need to project to to maintain pairwise distances is

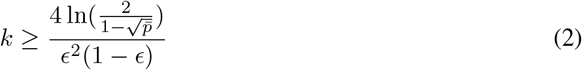

Substituting 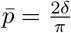 from the result in Section E.2, and *δ* = 2(1 − *p*) from Section E.1 into Eq. 2,

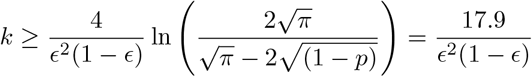

for *p* = 0.25 as mentioned in Section B.

Additionally, the above equation also implies that given *k* = 32 which is the smallest dimension we project to in our microcircuit, the distortion |*ϵ*| ≥ 0.592 for a normalized Bernoulli input. Likewise to minimize distortion of the inputs, we would want *k* ≈ 120.

While these results hold precisely for weights that are sampled from 𝒩 (0, 1), the nature of the results extends to other Gaussian distributions too, with changes in the constants. All weights at initialization in the microcircuit are sampled from the Gaussian distribution 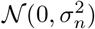 where *n* is the number of neurons and the variance 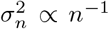. These include recurrent weights within a population (e.g., *W*_*LL*4_, *W*_*LL*5*/*6_, *W*_*HL*2*/*3_, etc.), weights of the inter-areal connections (i.e, *W*_*FF*_, *W*_*FBa*_, *W*_*FBb*_) and intra-areal lateral connections (i.e., *W*_*BB*_).

Furthermore, while we impose sparsity in the connectivity across and within layers by multiplying all non-recurrent unit weights with a binary mask sampled from a Bernoulli distribution ℬ (*p*), where *p* is a pre-determined probability of connection, the sparsified weights are simply a subset sampled from the original Gaussian distribution, and still follow a Gaussian distribution, viz. 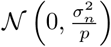.

In totality, the results and analyses in this section provide a mathematically grounded explanation as to why it is possible for the neuronal representations in *LL*2*/*3, *LL*5*/*6 to distinguish between expected and surprising elements of a sequence in a simple sequential memorization task. We note that these results require the tasks to be both “simple” and sequential in nature, as if the inputs are not significantly apart in the input/representation space before the lower cortical area, or if there is no explicit temporal structure that the microcircuit can exploit, this argument fails.

## F Performance across architectures and tasks in presence of expectation violations

**Figure 10:**
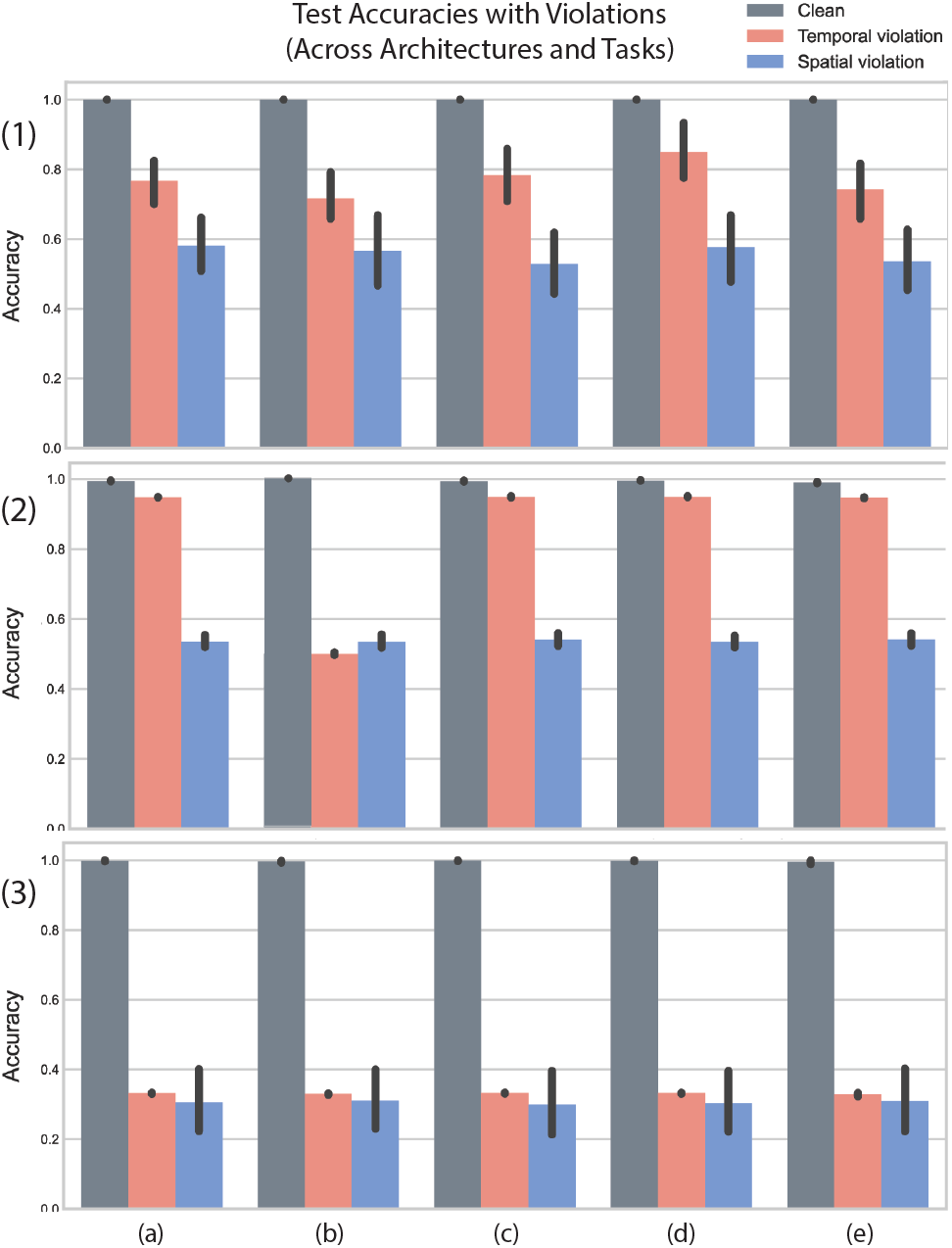
Performance of architectures across all tasks and violations. Rows denote different tasks and columns denote different architectural modifications of the cortical microcircuit. Task (1) = Sequence memorization, Task (2) = Binary Addition, Task (3) = Lattice Navigation. Architecture (a) = CorticalRNN, Architecture (b) = No Feedback, Architecture (c) = Bi-directional Feedback, Architecture (d) = Uni-directional Feedback, Architecture (e) = Population-controlled.

We note that the red bars (test accuracies for producing the correct final output given temporal violations in the input sequence) for architectures that receive feedback (columns a, c, d, e) are appreciably higher than that of the no-feedback architecture (column b) for the binary addition task (row 2) and slightly higher for the sequence memorization task (row 1).

## G Predictive coding based training algorithm

### Algorithm 1 Predictive-Coding Based Training

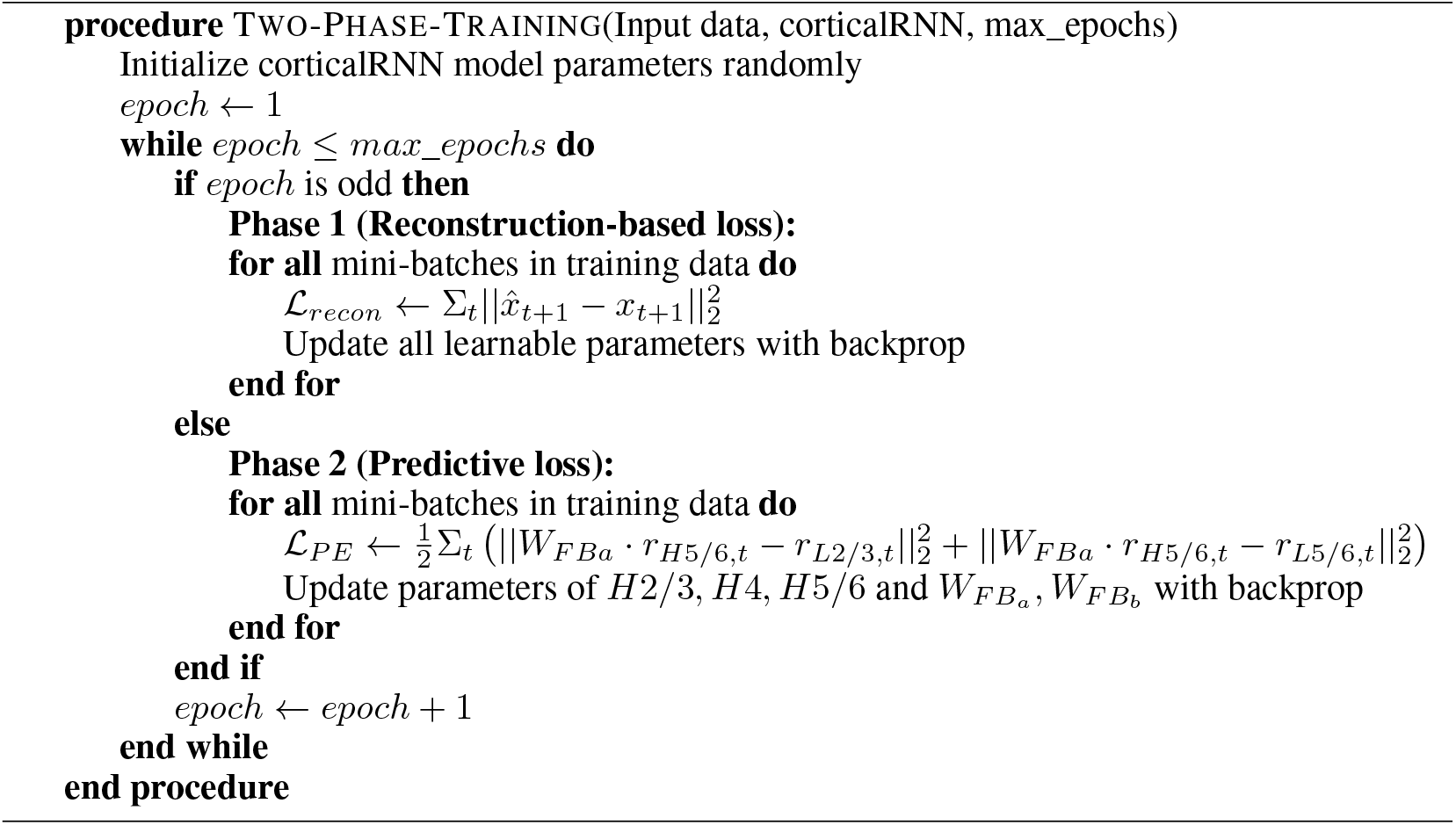

## H Dimensionality gain with predictive-coding (PC) loss across binary prediction and lattice navigation tasks

**Figure 11:**
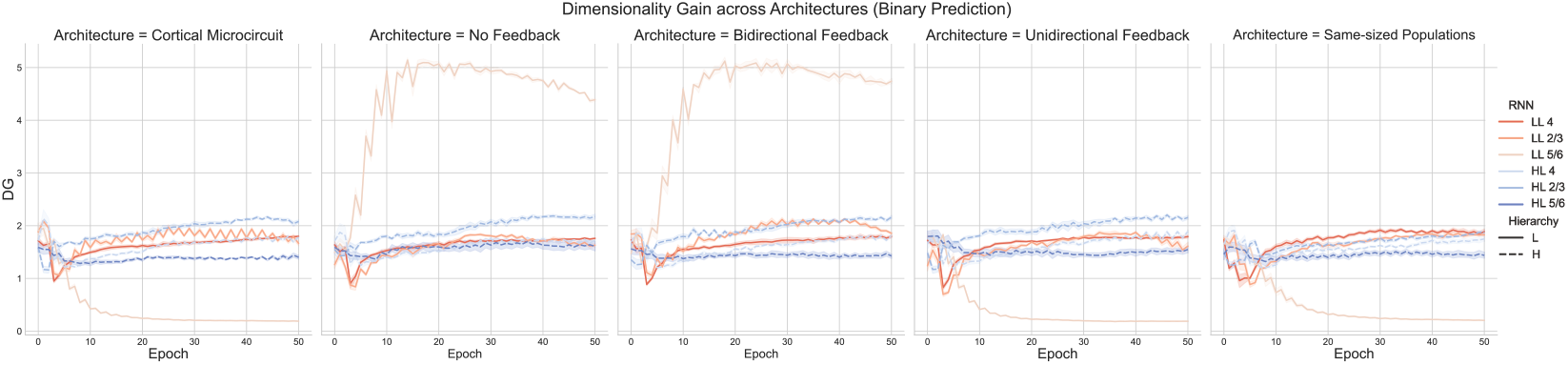
Binary prediction: Dimensionality gain across architectures

**Figure 12:**
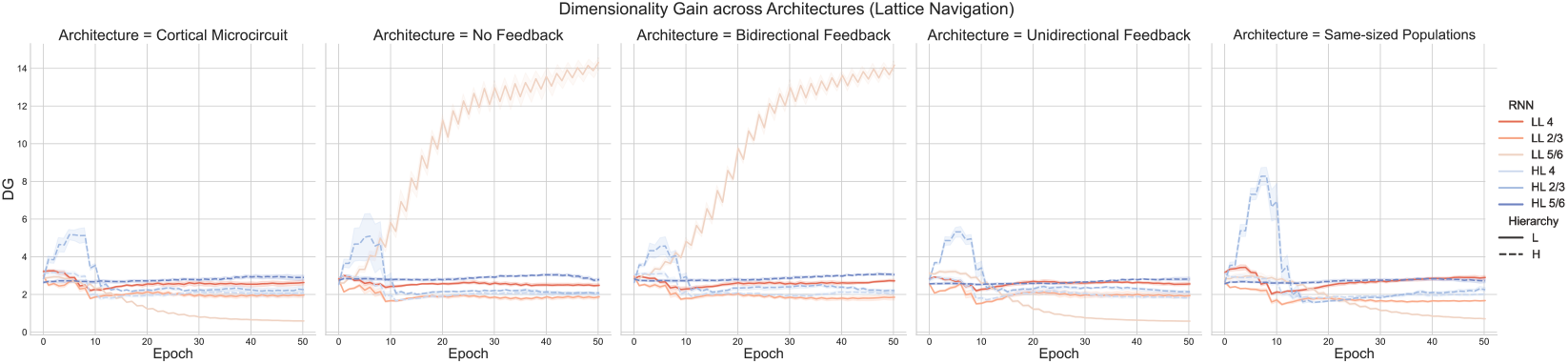
Lattice navigation: Dimensionality gain across architectures

## I Neuronal selectivity for surprise in LL5/6 with predictive-coding (PC) loss

**Figure 13:**
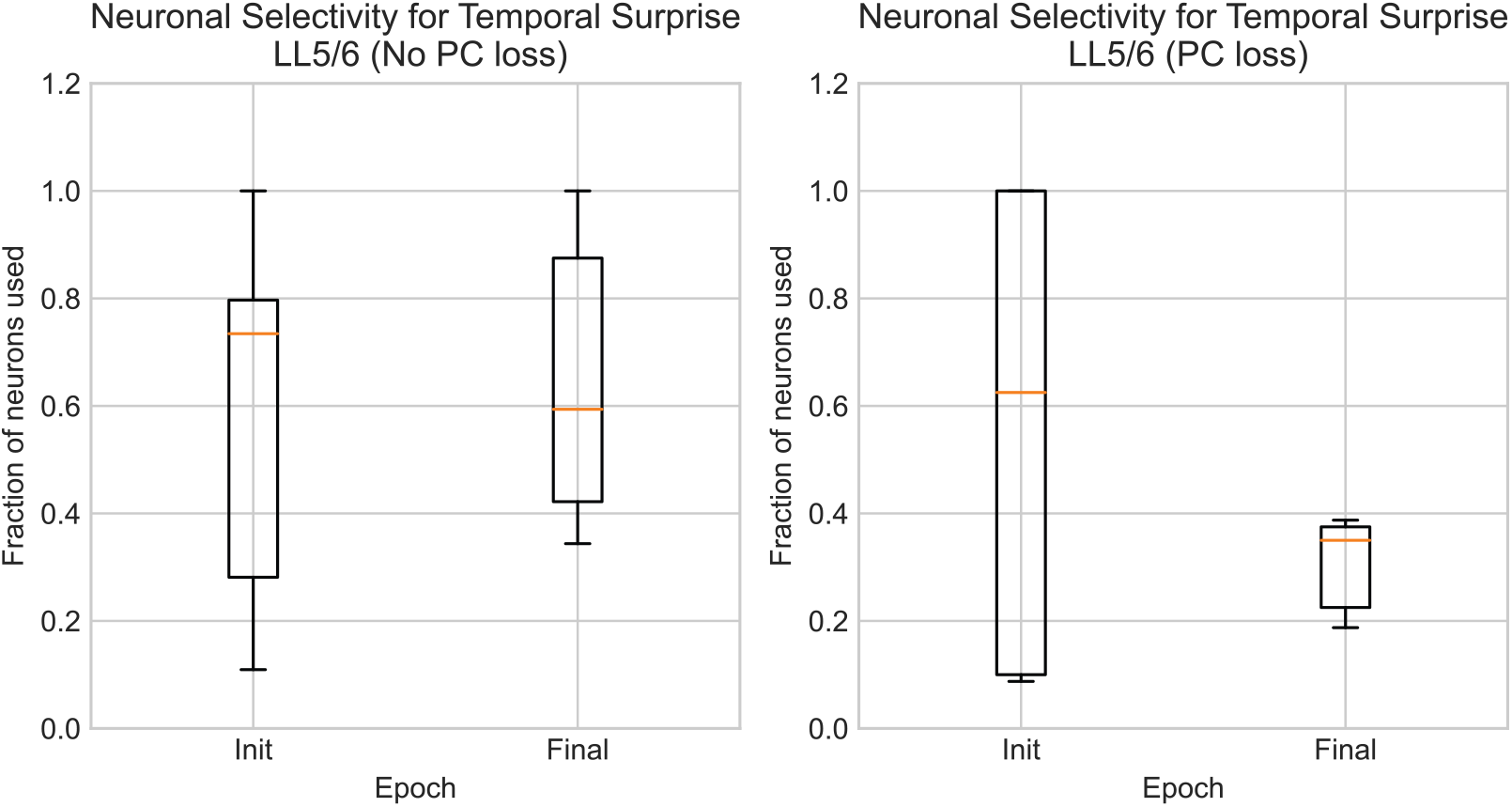
Effect of PC loss on neuronal selectivity for surprise in LL5/6. (Left) Fraction of neurons needed to decode surprise without PC loss at initialization and post training. (Right) Fraction of neurons needed to decode surprise with PC loss at initialization and post training.

We find that when averaged over multiple runs, the fraction of neurons needed to decode expected v/s surprise is consistently lower post-training with the PC loss compared to initialization, as well as when not using the PC loss. Moreover, the drop in the median fraction of neurons used is higher when the microcircuit is trained with the PC loss (Right) than without (Left).

The box and whisker plots follow standard convention, where the lower edge of the box extends to the first bottom (*Q*1) and the upper edge extends to the top of the third quartiles (*Q*3). The lower whisker extends to *Q*1 − 1.5* (*Q*3 − *Q*1) and the upper whisker extends to *Q*3 + 1.5* (*Q*3 − *Q*1). The line inside the box plots represents the median (*Q*2).

## Footnotes

4 LL 4 = Lower area layer 4, LL 5/6 = Lower area layers 5 & 6. Similar shorthand used for higher area layers, for example, HL 2/3 = Higher area layers 2 & 3

5 In the case where the the vectors *x, y* themselves are normalized to be unit length, the squared distance 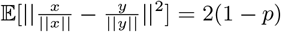 since 𝔼 [||*x*||^2^] = 𝔼 [||*y*||^2^] = *dp* acts as the common denominator and cancels out with part of the numerator for 𝔼 [||*x* − *y*||^2^] = 2*dp*(1 − *p*).

